# RAD52 and ERCC6L/PICH have a compensatory relationship for genome stability in mitosis

**DOI:** 10.1101/2023.08.23.554522

**Authors:** Beth Osia, Arianna Merkell, Felicia Wednesday Lopezcolorado, Xiaoli Ping, Jeremy M. Stark

## Abstract

The mammalian RAD52 protein is a DNA repair factor that has both strand annealing and recombination mediator activities, yet is dispensable for cell viability. To characterize genetic contexts that reveal dependence on RAD52 to sustain cell viability (i.e., synthetic lethal relationships), we performed genome-wide CRISPR knock-out screens. Subsequent secondary screening found that depletion of ERCC6L in RAD52-deficient cells causes reduced viability and elevated genome instability, measured as accumulation of 53BP1 into nuclear foci. Furthermore, loss of RAD52 causes elevated levels of anaphase ultrafine bridges marked by ERCC6L, and conversely depletion of ERCC6L causes elevated RAD52 foci both in prometaphase and interphase cells. These effects were enhanced with combination treatments using hydroxyurea and the topoisomerase IIα inhibitor ICRF-193, and the timing of these treatments are consistent with defects in addressing such stress in mitosis. Thus, loss of RAD52 appears to cause an increased reliance on ERCC6L in mitosis, and vice versa. Consistent with this notion, combined depletion of ERCC6L and disrupting G2/M progression via CDK1 inhibition causes a marked loss of viability in RAD52-deficient cells. We suggest that RAD52 and ERCC6L play compensatory roles in protecting genome stability in mitosis.

## INTRODUCTION

Human RAD52 (radiation sensitive 52) is a protein that self-associates to form an oligomeric ring structure and exhibits DNA-binding activities for both double-stranded and single-stranded DNA, which enables RAD52 to perform both annealing (Saotome et al. 2018; Jalan et al. 2019) and strand exchange functions (Jalan et al. 2019). RAD52 is also important for several types of homology-driven DNA repair, including single strand annealing (SSA) (Stark et al. 2004; Motycka et al. 2004; Rothenberg et al. 2008; Bhargava et al. 2016), homologous recombination (HR) associated with transcription (Yasuhara et al. 2018; Welty et al. 2018; Teng et al. 2018; Tan et al. 2020; Guha and Bhaumik 2022), and mitotic DNA synthesis (MiDAS) (Bhowmick et al. 2016; Min et al. 2017, 2019; Özer et al. 2018; Epum and Haber 2021). The latter (MiDAS) refers to synthesis that occurs in early mitosis (e.g., prophase) to repair under-replicated regions, which appears to be particularly important for replication of common fragile sites (Bhowmick et al. 2016; Özer et al. 2018; Minocherhomji et al. 2015; Di Marco et al. 2017). In addition to its roles in these aspects of HR, RAD52 is involved in alternative lengthening of telomeres (ALT) (Zhang et al. 2019; Verma et al. 2019) and plays a protective role at replication forks by preventing unscheduled fork reversal and degradation (Malacaria et al. 2019).

While RAD52 appears to be involved in several aspects of genome maintenance, the circumstances that require such functions of RAD52 remain poorly understood, particularly since RAD52 is not essential for viability (Rijkers et al. 1998). One approach to identifying such circumstances is to evaluate synthetic lethal relationships with RAD52. Synthetic lethality is generally defined as the disruption of two cellular factors resulting in reduction of viability or proliferation, while perturbation of either factor individually does not have such an effect (Patel et al. 2021; Akimov and Aittokallio 2021) RAD52 has been reported to show synthetic lethal relationships with several HR factors (e.g. BRCA1, BRCA2, PALB2, and RAD51 paralogs) (Jalan et al. 2019; Patel et al. 2021; Hengel et al. 2017; Lok et al. 2013; Feng et al. 2011; Chun et al. 2013). Beyond such HR deficiencies, the range of synthetic lethal interactions with RAD52 is largely undefined. Given that numerous RAD52 small molecule inhibitors have been described that may be translatable for use in the clinic (Sullivan-Reed et al. 2018; Hengel et al. 2017; Patel et al. 2021), expanding the understanding of when cells become reliant on RAD52 will inform the successful application of such inhibitors in cancer treatment.

Here, we describe a set of synthetic lethality screens with RAD52 loss, and from secondary screening describe a set of studies with ERCC6L (also known as PICH). ERCC6L is a SWI-SNF2-family DNA translocase involved in the resolution of DNA bridges that form between sister chromatids in mitosis due to incomplete replication (Chan et al. 2009, 2018; Joseph et al. 2020; Baumann et al. 2007; Chan et al. 2007). ERCC6L has also been reported to largely promote genome stability during mitosis, and is excluded from the nucleus prior to mitotic nuclear envelope breakdown (Baumann et al. 2007; Kaulich et al. 2012). In mitosis, ERCC6L localizes to ultrafine DNA bridges (UFBs), which are non-chromatinized DNA bridges that form at points of sister chromatid nondisjunction as cells prepare to enter mitotic anaphase (Chan et al. 2007). Evidence supports that ERCC6L initiates the dissolution of UFBs in a process involving recruitment of BLM, topoisomerase IIIα (TopoIIIα), RMI1, and RMI2 (together known as the BTRR complex), and topoisomerase IIα (TopoIIα) (Baumann et al. 2007; Chan et al. 2007; Kaulich et al. 2012; Nielsen et al. 2015; Sarlós et al. 2018; Ke et al. 2011; Rouzeau et al. 2012). Biochemical analysis of ERCC6L suggests that the protein recognizes and binds to DNA under tension (e.g., that of DNA catenanes as sister chromatids segregate during anaphase), and that ERCC6L, together with TopoIIIα, produces positive supercoiling required for rapid decatenation of sister chromatids by TopoIIα at anaphase onset (Biebricher et al. 2013; Bizard et al. 2019). Given that RAD52 and ERCC6L are both implicated in genome maintenance in mitosis, and because we identified ERCC6L in our synthetic lethal screen with RAD52, we sought to examine the interplay between these two factors for genome stability, lethality, and the influence of RAD52 on formation of ERCC6L-UFBs, and conversely the influence of ERCC6L on localization of RAD52 into foci.

## RESULTS

### Genome-wide CRISPR knockout screen identifies pathways that are synthetic lethal with RAD52

To identify genes and pathways that are synthetic lethal with RAD52, we performed genome-wide CRISPR-Cas9 knockout screens to measure loss of fitness for genes knocked out concurrently with RAD52. Namely, we sought to identify genes that when knocked out (KO) cause reduced fitness specifically in RAD52^KO^ cells as compared to RAD52^WT^ cells, which we refer to as the “synthetic lethal hits.” To this end, we generated a RAD52^KO^ cell line from a cell line used previously for DNA damage response screens (RPE-1 hTERT p53^KO^ Cas9) (Zimmermann et al. 2018; Olivieri et al. 2020; Olivieri and Durocher 2021). The parental cell line is RPE-1, which is a human epithelial cell line immortalized with hTERT. We used a version of this cell line that is p53^KO^, which facilitates survival of cells that undergo CRISPR-Cas9 gene editing (Haapaniemi et al. 2018), and because loss of p53 is relevant to many cancers (Kandoth et al. 2013). This line also stably expresses Cas9, such that only the sgRNA component is transduced during the screen. We refer to this cell line as the RAD52^WT^ line. From this cell line, we generated the RAD52^KO^ line by Cas9-mediated deletion of a region between exons 3 and 4 of RAD52 using two previously described sgRNAs, resulting in loss of the RAD52 protein (Kelso et al. 2019) (Fig 1a).

**Figure 1:**
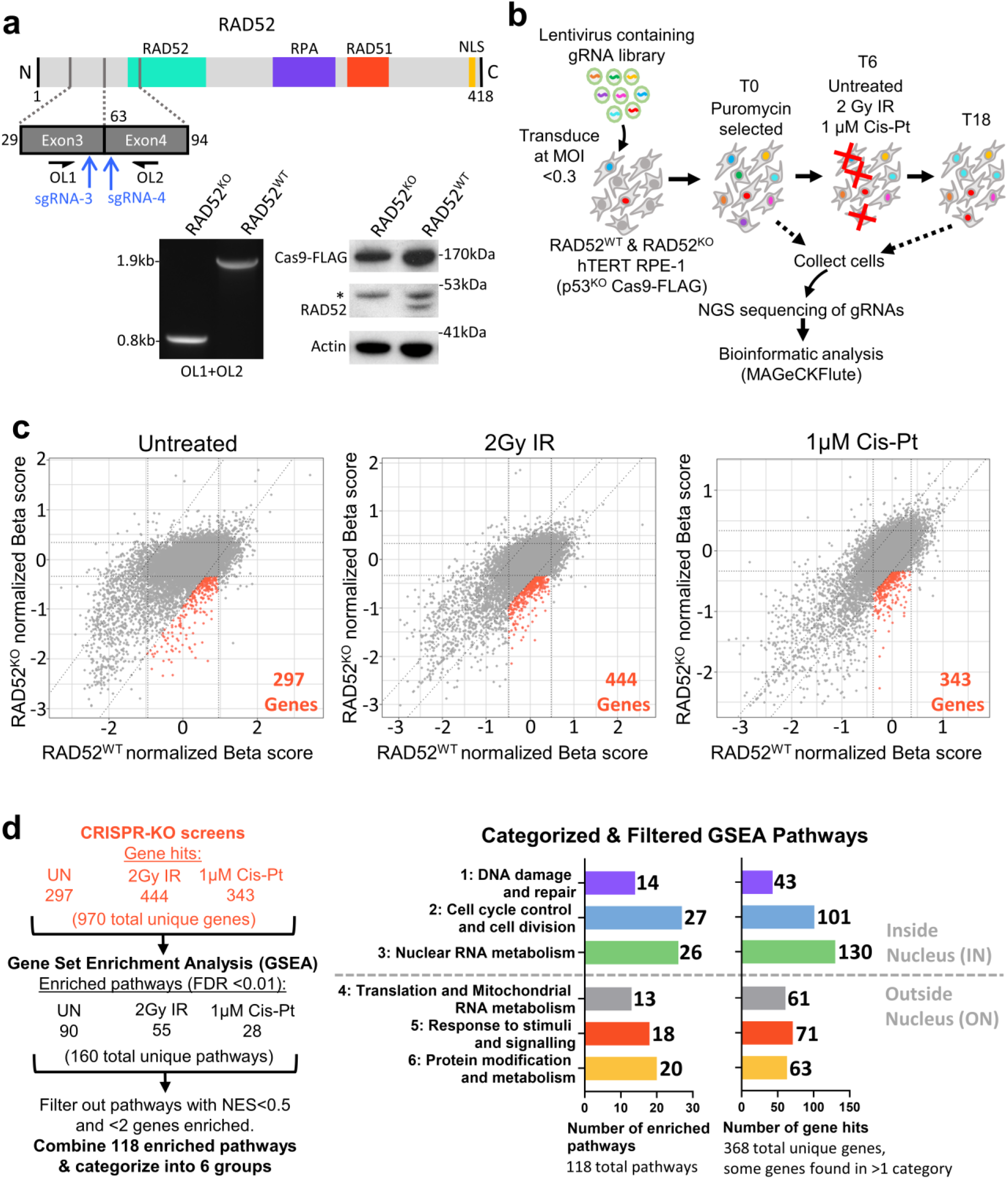
Genome-wide CRIPSR knockout screen identifies pathways that are synthetic lethal with RAD52. **a**) The RAD52 knockout (RAD52^KO^) cell line using p53^KO^ Cas9-FLAG-expressing RPE-1 hTERT (RAD52^WT^). Shown is a schematic of RAD52 with blue arrows depicting two sgRNAs used to generate a deletion in the *RAD52* gene, which was assessed by PCR using primers OL1 and OL2 (left gel), along with western blot (right blot). *non-specific band. **b)** Schematic of genome-wide CRISPR knockout screen to identify RAD52 synthetic lethal interactors. **c)** Shown are synthetic lethal hits (orange data points) identified under each screening condition by applying cutoffs of 1.5x the standard deviation from the diagonal axis, x=0 and y=0 of the Beta (selectivity) scores generated by MAGeCKFlute. **d)** Enriched cellular pathways from RAD52 synthetic lethal hits identified in (c) identified 6 groups of pathways. Shown is the analysis pipeline (left) and the 6 groups of pathways that are enriched among screen hits from all conditions (right). Groups 1-3 represent pathways that function inside the nucleus (IN) and groups 4-6 represent pathways that function outside the nucleus (ON).

With these two cell lines, we performed a set of genome-wide CRISPR knockout screens to compare gene knockouts that affect the fitness of this RAD52^KO^ line as compared to its parental RAD52^WT^ line. We performed the screens by transducing the RAD52^KO^ and RAD52^WT^ cell lines with lentivirus containing a genome wide knockout CRISPR library (Addgene Pooled Library #67989) (Tzelepis et al. 2016) at low multiplicity of infection (<0.3 MOI). Genomic DNA was prepared from a timepoint shortly after selection of transduced cells (set as time day 0 / T0), which is the reference sample for the screen. Subsequently, the cells were split into 3 treatment groups: 1) cells left untreated for 18 days after T0 (T18), 2) cells exposed to 2 Gy ionizing radiation (IR) on day T6, and then cultured for 12 days (total T18), and 3) cells treated with 1 µM Cisplatin (Cis-Pt) at T6 and for 6 days (T12), and then cultured for 6 more days (total T18) (Fig 1b). We performed three different treatment conditions to broaden the scope of our overall screen.

Genomic DNA was then prepared at T18 for each cell line and exposure condition, along with the T0 sample for each cell line, and guide sequences were amplified and sequenced using the Illumina HiSeq platform. Sequencing results were analyzed using the maximum likelihood estimation (MLE) algorithm from the MAGeCK analysis pipeline (Li et al. 2014), which generates selectivity values (known as beta-scores) by comparing guide counts from the T18 timepoint to the T0 timepoint for each gene represented in the library (Fig 1b, Supplementary Table 2). We then applied the beta-score normalization method from the MAGeCKFlute analysis suite, which uses each sample’s beta-scores and a list of common essential genes to normalize all beta-scores between samples to account for cell division rate differences (Wang et al. 2019). We plotted the normalized beta-scores for all genes from the RAD52^WT^ screens (x-axis) against the RAD52^KO^ screens (y-axis) (Fig 1c). We applied cutoffs of 1.5-fold from the standard deviation to the x, y, and diagonal axes. From these cutoffs, we determined the “synthetic lethal hits” to be genes that are negatively selected in the RAD52^KO^ line, but non-selected in the RAD52^WT^ line (Fig 1c, orange sections). These “synthetic lethal hits” amounted to 297, 444, and 343 genes for the untreated, IR-treated, and Cis-Pt-treated screens respectively (Fig 1c-d, Supplementary Table 2). When combined, these screens produced 970 unique genes (i.e., genes not repeated between screens) (Fig 1d).

To identify a set of genes for secondary screening, we performed gene set enrichment analysis (GSEA) on the gene hits from each of the three screens separately (Fig 1d). The goal of this analysis approach was to identify cellular pathways and processes important for cellular fitness with RAD52 loss. We queried multiple functional and pathway databases (e.g., Gene Ontology, Reactome, Kyoto Encyclopedia of Genes and Genomes) using the GSEA package included with MAGeCKFlute and imposed a cut off such that the false discovery rate (FDR) for each pathway at < 0.01. In doing so, we identified 90 (untreated), 55 (IR), and 28 (Cis-Pt) enriched pathways, with 160 unique (unrepeated) pathways between all three screens (Fig 1d, Supplementary table 3). We then filtered out any pathways with fewer than 2 enriched genes or a normalized enrichment score (NES) of 0.5 or less, where a high NES indicates that enriched gene hits are distributed within known functional roles (as opposed to randomly) in a pathway (Fig 1d). This filtering led to 118 pathways total (Fig 1d).

We combined these 118 filtered pathways and categorized them into 6 groups: (1) DNA damage and repair (14 pathways and 43 genes), (2) Cell cycle control and cell division (27 pathways and 101 genes), (3) Nuclear RNA metabolism (26 pathways and 130 genes), (4) Translation and Mitochondrial RNA metabolism (13 pathways and 61 genes), (5) Response to stimuli and signaling (18 pathways and 71 genes), and (6) Protein modification and metabolism (20 pathways and 63 genes) (Fig 1d, Supplementary table 3). We then focused on the first three groups, which involved processes that function inside the nucleus (IN) vs. those function outside the nucleus (ON), because of the established role of RAD52 in DNA Repair(Bhat et al. 2022) (Fig 1d). Furthermore, IN pathways made up the greatest proportion of enriched pathways (67 IN vs. 51 ON pathways). Finally, to narrow down the screen hits identified by GSEA for secondary screening, we selected 59 gene hits from those enriched IN pathways (i.e., 59 / 232) (Supplementary Table 3). These genes were selected to cover all 67 pathways from the three IN pathway groups, with at least one gene selected from each pathway, and some genes participating in multiple pathways (Fig 2a, Supplementary table 3).

**Figure 2:**
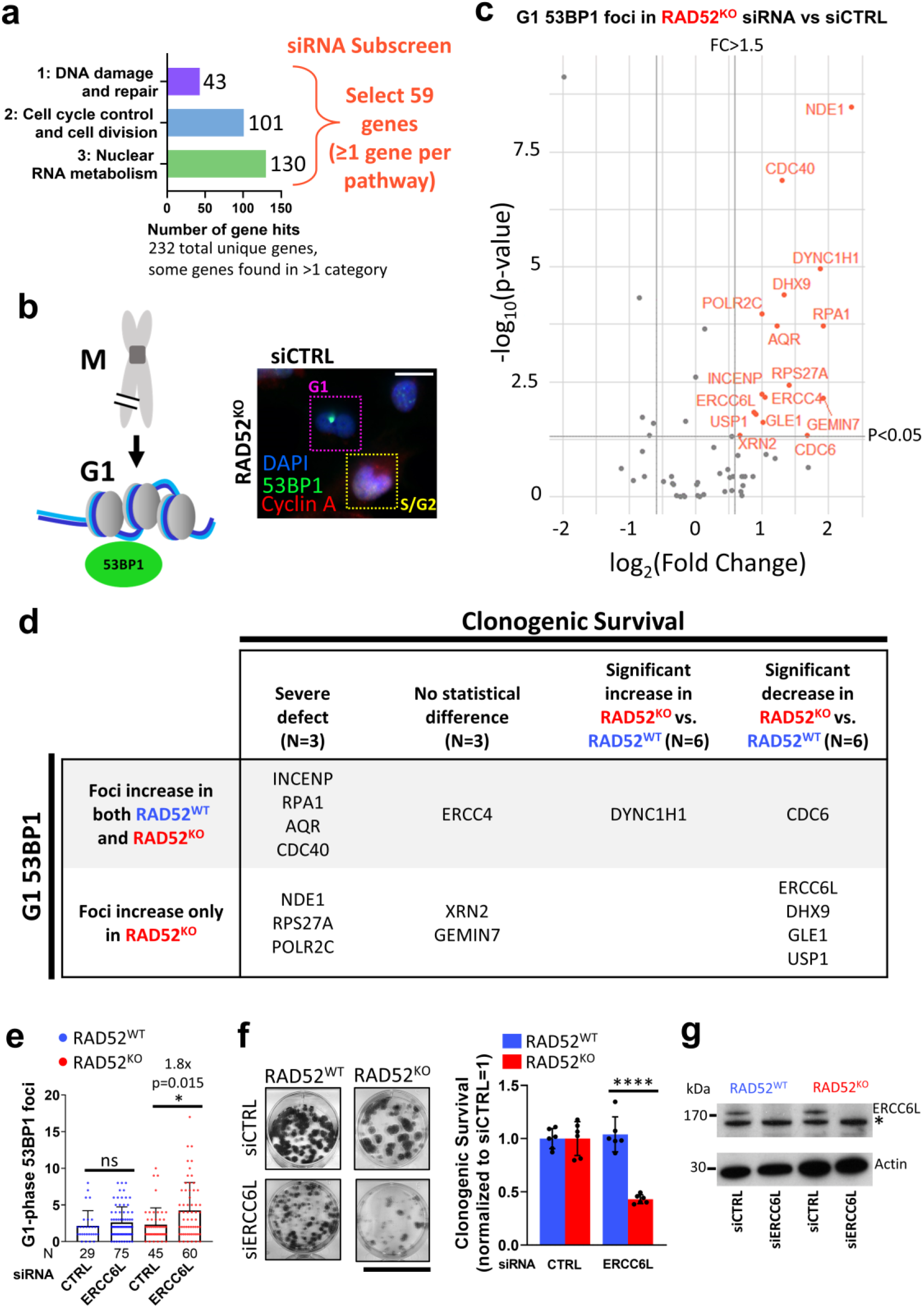
Secondary screen of 59 genes identifies 5, including ERCC6L/PICH, for which depletion causes G1-phase 53BP1 foci and reduced clonogenic survival in RAD52^KO^ cells. **a**) Strategy for selecting 59 genes from IN Groups 1-3 (see Fig 1e) for secondary screening. **b)** 53BP1 foci assay. Shown is an illustration that DSBs that persist through mitosis (M) form G1 53BP1 nuclear bodies (left), along with a representative immunofluorescence image of 53BP1 and Cyclin A. Scale bar is 20 µm. Image taken at 20x magnification. **c)** Effects of siRNAs targeting the 59 genes in (a) on G1 53BP1 foci in RAD52^KO^ cells, compared to non-targeting control siRNA (siCTRL). Highlighted in red are the siRNAs that caused a significant, i.e., p<0.05 by Kolmogorov-Smirnov (K-S) test and >1.5-fold mean, increase in G1 53BP1 foci. N>50 total nuclei (G1 and S/G2) per siRNA. **d)** Shown are summary data for the genes in red shown in (c) for effects on 53BP1 G1 foci, as well as clonogenic survival, in RAD52^KO^ and not in RAD52^WT^ cells. Effects on 53BP1 G1 foci determined as in (c), and significant difference in clonogenic survival based on unpaired *t*-test (p<0.05). **e)** G1 53BP1 foci increase with siERCC6L treatment in RAD52^KO^, but not RAD52^WT^ cells. ns=not significant, *=p<0.05 by K-S test, bars show mean foci value. Mean fold-increase and K-S test p-value are shown for significant comparisons. N (number of cells) is shown below each bar. **f)** Clonogenic survival is reduced in RAD52^KO^ cells with ERCC6L depletion. Colonies from representative wells are shown (left), along with clonogenic survival normalized to the respective siCTRL treated lines (siCTRL=1) (right). Scale bar = 35mm, ns=not significant, *=p<0.05 by unpaired t-test. N=6 replicates. **g)** Western blot confirming ERCC6L depletion via siERCC6L in RAD52^KO^ and RAD52^WT^ cell lines. *non-specific band.

### Secondary screen of 59 genes identifies 5, including ERCC6L/PICH, for which depletion causes G1-phase 53BP1 foci and reduced clonogenic survival in RAD52^KO^ cells

Given the role of RAD52 in DNA repair, we hypothesized that some of the synthetic lethal hits may be involved in mitigating genotoxic stress, particularly when RAD52 is absent. One method to assay for genotoxic stress is to measure 53BP1 accumulation into foci, which occurs as part of the DNA damage response (DDR) (Mirza-Aghazadeh-Attari et al. 2019). More specifically, 53BP1 foci that form during G1-phase of the cell cycle (also known as G1 53BP1 nuclear bodies) appear to mark chromosomal breaks that originated from replication stress in the prior cell cycle that persisted through mitosis (Lukas et al. 2011b, 2011a; Harrigan et al. 2011) (Fig 2b). Furthermore, a fraction of such G1 53BP1 foci persist into the subsequent S-phase where they are dissolved, presumably as damage is resolved to prevent further instability (Spies et al. 2019). Thus, we sought to use G1 53BP1 foci as an indicator of genotoxic stress that persists through mitosis. Meanwhile, we also sought to assess 53BP1 foci later in the cell cycle (S/G2-phase), which might indicate passage of such genotoxic lesions into S-phase, or response to new S-phase lesions.

Thus for secondary screening, we depleted each of the 59 of the selected genes described above (Fig 2a) in RAD52^KO^ and RAD52^WT^ cells by siRNA (pools of 4 siRNAs per gene) and examined 53BP1 foci by immunofluorescence microscopy. We also co-stained cells for Cyclin A to differentiate between S/G2-phase cells (Cyclin A positive) and G1-phase (Cyclin A negative) (Lukas et al. 2011a; Harrigan et al. 2011) (Fig 2b). Additionally, we included control cells that were treated with non-targeting siRNA (siCTRL). From the 59 genes analyzed (Supplementary Table 4), depletion of 16 produced significantly increased levels of G1 53BP1 foci (i.e., p<0.05, and a greater than 1.5-fold increase of the mean number of foci vs. siCTRL), in either RAD52^KO^ cells alone, or both RAD52^KO^ and RAD52^WT^ cells (Fig 2c, Supplementary Fig 1a-b). Additionally, 17 of the 59 genes produced significantly increased levels of S/G2 53BP1 levels when depleted in the RAD52^KO^ or both lines (Supplementary Fig 1c-d). Among these genes, 8 showed an increase in 53BP1 foci in both G1 and S/G2 (Supplementary Fig 1b, d). However, we chose to focus on the 16 genes for which depletion caused an increase in G1 53BP1 foci for two reasons: 1) such foci appear to be an indicator of damage transmitted through mitosis (Lukas et al. 2011a; Harrigan et al. 2011; Barr et al. 2017) (although spontaneous lesions generated in G1 are also possible), and 2) RAD52 has been shown to have a role in DNA repair in mitosis, particularly for promoting mitotic DNA synthesis (MiDAS) (Bhowmick et al. 2016; Min et al. 2017, 2019; Özer et al. 2018; Epum and Haber 2021).

Subsequently, we assessed effects of siRNAs targeting these 16 genes on viability in RAD52^KO^ cells and RAD52^WT^ by a clonogenic survival assay (normalized to parallel siCTRL treatment in each cell line, supplementary Fig 2a). We found that siRNAs targeting 5 genes caused a loss of viability that was significantly more severe in the RAD52^KO^ line as compared to the RAD52^WT^ line (Fig 2d). The remaining 11 genes tested by siRNA either showed severe viability defects in both cell lines (7 genes), no significant difference between the two lines (3 genes), or improved viability in the RAD52^KO^ line compared to the RAD52^WT^ line (1 gene) (Fig 2d). Of the 5 genes with viability defects in the RAD52^KO^ line, siRNA targeting of 4 showed a specific increase in G1 53BP1 foci in RAD52^KO^ but not RAD52^WT^ cells (i.e., ERCC6L, DHX9, GLE1, and USP1), whereas siRNA targeting of the fifth gene (CDC6) caused an increase in G1 53BP1 foci in both lines (Fig 2d).

As an example, ERCC6L was identified by the IR-treated genome-wide screen, through GSEA was categorized in IN group 2, and was selected as one of the 59 genes for secondary screening (Supplementary Table 3). From the secondary screening we found that siRNA targeting ERCC6L (siERCC6L) caused a significant (1.8-fold) increase in the mean number of G1 53BP1 foci compared to siCTRL in RAD52^KO^ cells, while there was no significant difference between ERCC6L depletion vs. siCTRL in RAD52^WT^ cells (Fig 2e). Moreover, siERCC6L treatment produced a significantly greater loss of clonogenic survival in RAD52^KO^ cells vs. RAD52^WT^ cells (Fig 2f). Lastly, we confirmed siRNA depletion of the ERCC6L protein by western blot analysis in both the RAD52^KO^ and RAD52^WT^ cell lines (Fig 2g).

### RAD52 suppresses the accumulation of ERCC6L Ultra-Fine DNA bridges in anaphase

We then sought to further characterize the interrelationship between RAD52 and ERCC6L. ERCC6L (also known as PICH) is a SWI/SNF2-family ATPase that interacts with the mitotic kinase PLK1, and has been shown to be critical for the resolution of DNA ultrafine bridges (UFBs) that form during anaphase of mitosis at centromeres and late-replicating regions or as a result of replication stress (Chan et al. 2009; Özer and Hickson 2018; Chan and Hickson 2011). Furthermore, ERCC6L itself localizes to such UFBs (i.e., ERCC6L-UFBs) (Chan et al. 2007; Baumann et al. 2007). To begin to understand how ERCC6L might compensate for the loss of RAD52 in mitigating persistent genotoxic stress, we first tested whether the loss of RAD52 would cause an increase in ERCC6L-UFBs. Namely, we posited that without RAD52, resolution of chromosomes in anaphase may be more reliant on ERCC6L-coated UFBs, as assessed by an increased frequency of these structures. To test this hypothesis, we performed immunofluorescence (IF) staining of the ERCC6L protein in RAD52^KO^ and RAD52^WT^ cells and quantified ERCC6L-coated UFBs in anaphase cells (Fig 3a-b). From this analysis, levels of ERCC6L-coated UFBs per anaphase were significantly increased in the RAD52^KO^ line as compared to the RAD52^WT^ line (Fig 3b).

**Figure 3:**
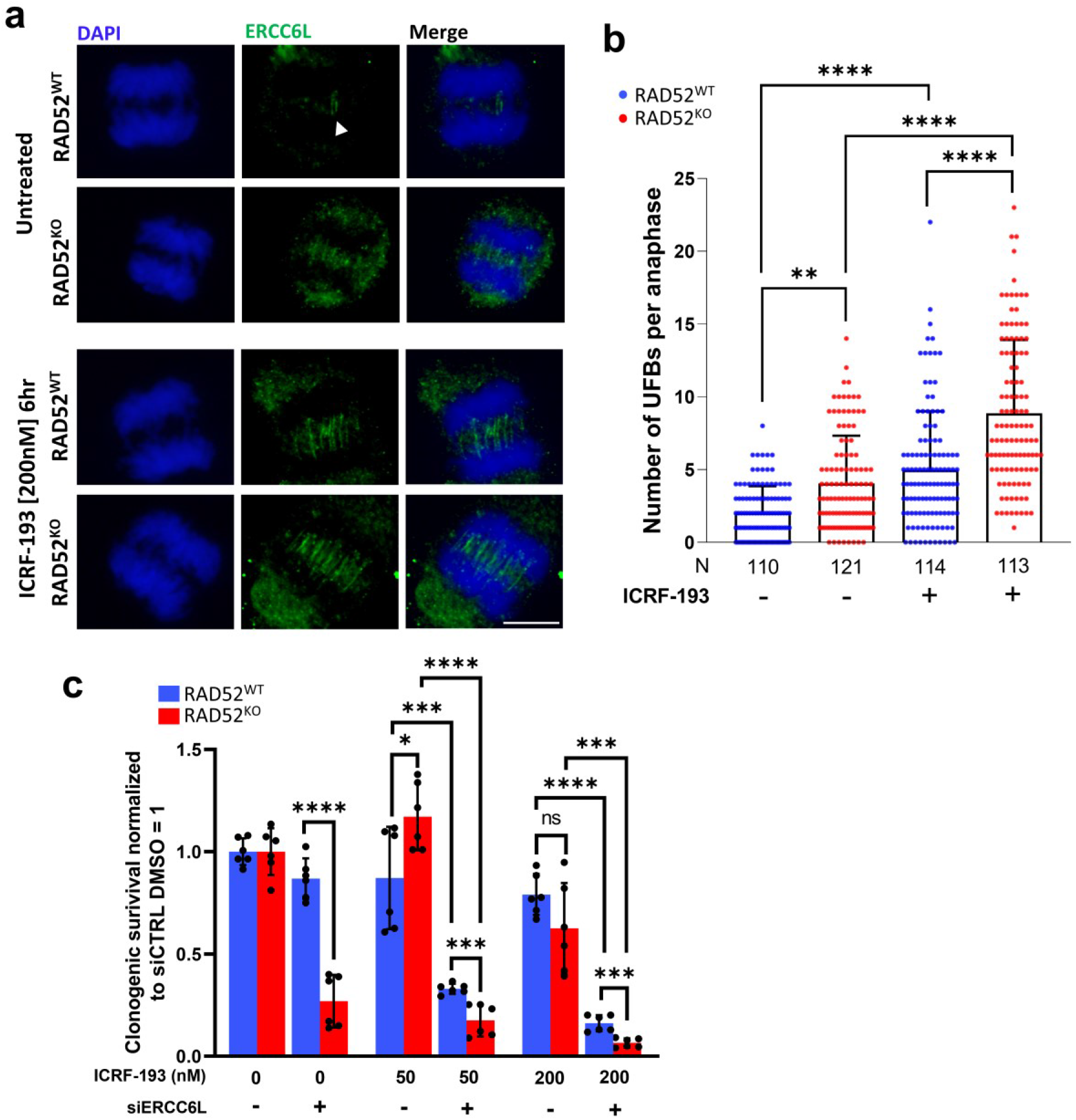
RAD52 suppresses the accumulation of ERCC6L Ultra-Fine DNA bridges in anaphase. **a**) ERCC6L UFBs. Shown are examples of anaphase cells stained with ERCC6L for RAD52^WT^ and RAD52^KO^ cells with and without exposure to ICRF-193 [200mM, 6-hour treatment]. White arrowhead indicates ERCC6L-UFBs in untreated RAD52^WT^ cells. Scale bar is 10µm, and images were taken at 60x magnification. **b)** Anaphase ERCC6L-UFBs are increased in RAD52^KO^ cells as compared to RAD52^WT^ cells, which is amplified by exposure to ICRF-193 as shown in (a). Bars show mean value. N (number of anaphases) is shown below each bar. Significance determined by K-S test, **=p<0.01, ****=p<0.0001. **c)** ICRF-193 exposure significantly impacts viability in ERCC6L-depleted RAD52^WT^ and RAD52^KO^ cells but does not significantly impact viability in RAD52^KO^ cells without ERCC6L depletion. ICRF-193 treatments were for 55-hours. ns=not significant, *=p<0.05, ***=p<0.001, and ****=p<0.0001. unpaired *t*-test. N=6 replicates.

Another factor important for mitigating UFBs is Topoisomerase IIα (Topo IIα), in that inhibition of Topo IIα by ICRF-193 increases the frequency of ERCC6L-UFBs (Baumann et al. 2007; Hengeveld et al. 2015; Gemble et al. 2020). Thus, we tested whether loss of RAD52 may also cause an increase in ERCC6L-UFBs caused by Topo IIα inhibition. For this, we treated cells with 200 nM ICRF-193 for 6 hours prior to fixation and IF staining, and we found that ICRF-193 treatment significantly increased ERCC6L-coated UFBs in both the RAD52^KO^ and RAD52^WT^ lines as compared to their untreated counterparts, however the most severe effect was observed in the RAD52^KO^ line (Fig 3a-b). These findings indicate that loss of RAD52 increases the frequency of ERCC6L-coated UFBs and chromatin bridges both under unstressed/spontaneous conditions, and with Topo IIα inhibition.

We then tested the effects of combined depletion of ERCC6L and ICRF-193 treatment on clonogenic survival in RAD52^KO^ cells. We treated RAD52^KO^ and RAD52^WT^ cells with 50 nM or 200 nM ICRF-193 after transfection with siERCC6L or siCTRL (i.e., a 2-day ICRF-193 treatment, which was then removed, and followed by assessment of colony formation). For both concentrations of ICRF-193, RAD52^KO^ cells did not show a significant decrease in clonogenic survival vs. RAD52^WT^ cells (treated with siCTRL) (Fig 3c). In contrast, a combination treatment of siERCC6L with ICRF-193 caused a decrease in clonogenic survival with both RAD52^KO^ and RAD52^WT^ cells, but with a greater decrease in RAD52^KO^ cells (Fig 3c). This marked loss of viability was most severe in RAD52KO cells treated with 200nM ICRF-193 and siERCC6L (Fig 3c). These findings indicate that RAD52 is important for survival following disruption of two factors important for resolving replication stress at mitosis (i.e., Topo IIα and ERCC6L).

### RAD52 forms foci in prometaphase cells in response to ERCC6L depletion and HU treatment, but the latter is dependent on G2/M arrest

Given that loss of RAD52 appears to cause an increased reliance on ERCC6L-UFBs in anaphase, we next sought to test the converse hypothesis, i.e., that cells may rely more heavily on RAD52 when ERCC6L is depleted, particularly in mitosis. To test this, we stably transfected hTERT RPE-1 P53^KO^ cells with GFP-tagged RAD52 (RAD52-GFP) to examine RAD52 focal accumulation, as previously described (Llorens-Agost et al. 2021). We treated these cells with siERCC6L vs. siCTRL, and then exposed cells to replication stress via treatment with hydroxyurea (HU) that depletes nucleotide pools (4-hour treatment with 2 mM HU, followed by 12-hour recovery). We also performed a mock treatment without HU (DMSO control). Following this treatment, we enriched for mitotic cells by treating cells with the CDK1 inhibitor RO3306 at a concentration of 7 µM for 6 hours to synchronize cells in G2/M, and then released cells into mitosis. 30 minutes after the release, cells were fixed, and we assessed RAD52-GFP foci by IF microscopy in prometaphase cells (Fig 4a). We observed marked increases in RAD52-GFP foci in siERCC6L treated cells as compared to siCTRL treated cells both with and without HU exposure (Fig 4b-c). Additionally, cells treated with HU produced significantly increased RAD52-GFP foci levels as compared to their untreated counterparts (Fig 4c). These results indicate that RAD52 localizes to prometaphase cells in response to ERCC6L depletion and HU-induced replication stress, with the greatest increase caused by the combined treatment.

**Figure 4:**
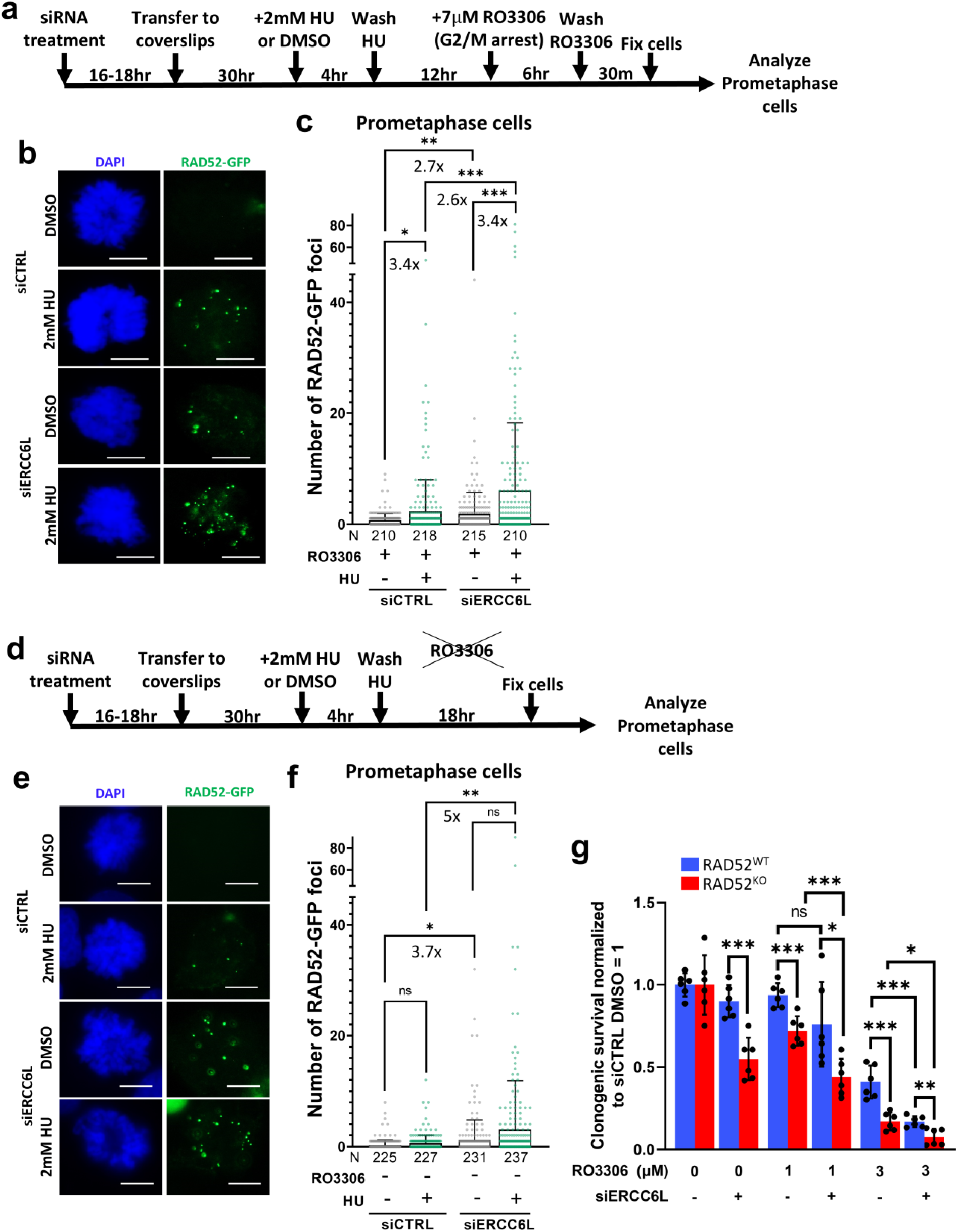
RAD52 forms foci in prometaphase cells in response to ERCC6L depletion and HU treatment, but the latter is dependent on G2/M arrest. **a**) Schematic of treatments used to examine RAD52-GFP foci in prometaphase cells. **b)** Shown are representative images of prometaphase cells with RAD52-GFP foci, after treatments as shown in (a). Scale bars are 10 µm, and images were taken at 40x magnification. **c)** Treatments with siERCC6L and HU each induce significant increases in RAD52-GFP foci in prometaphase cells. Cells were treated as in (a) and analyzed as in (b). Shown are mean foci values, along with fold increases. *=p<0.05, **=p<0.01, and ***=p<0.001, K-S test. N (number of cells) is shown below each bar. **d)** Schematic of treatments to examine RAD52-GFP foci without using the CDK1 inhibitor RO3306 (i.e., no induction of G2/M arrest). **e)** Shown are representative images of RAD52-GFP foci from the treatments as shown in (d). Scale bars are 10 µm and images were taken at 40x magnification. **f)** Without RO3306-induced G2/M arrest, RAD52-GFP foci are increased with siERCC6L treatment, but not HU treatment. Cells were treated as in (d) and analyzed as in (e). Shown are mean foci values and fold increases. N is shown below each bar. ns=not significant, *=p<0.05, **=p<0.01, K-S test. **g)** RO3306 exposure significantly impacts survival in RAD52^KO^ cells with and without ERCC6L depletion, and ERCC6L-depleted RAD52^WT^ cells at the high (3µM), but not low (1µM) concentration. Clonogenic survival assay was performed with two concentrations of RO3306 (1µM or 3µM) for 48 hours in RAD52^WT^ and RAD52^KO^ cell lines. Clonogenic survival is normalized to DMSO and siCTRL treated RAD52^WT^ and RAD52^KO^ cell lines (DMSO siCTRL = 1). ns=not significant, *=p<0.05, **=p<0.01, ***=p<0.001, unpaired *t*-test. N=6 replicates.

While RO3306 treatment is a common approach to enrich prometaphase cells, recent studies have indicated that such treatment can affect the cellular response to replication stress(Brison et al. 2023). Thus, we wondered if RO3306 treatment may influence the effect of ERCC6L depletion and/or HU treatment on RAD52-GFP foci formation. Thus, we assessed RAD52-GFP foci in cells treated with siERCC6L, with and without HU treatment, but without the addition of RO3306 (Fig 4d). We quantified RAD52-GFP foci in prometaphase cells and found that siERCC6L treatment caused a significant increase in RAD52-GFP foci as compared to siCTRL (Fig 4e-f), similar to the experiments with RO3306 treatment (Fig 4b-c). In contrast, RAD52-GFP foci were not significantly increased with the addition of HU as compared to untreated cells, both for siCTRL and siERCC6L treated cells (Fig 4f). Thus, ERCC6L depletion causes RAD52-GFP foci in prometaphase cells irrespective of induction of G2/M arrest via RO3306, whereas HU only induces such RAD52-GFP foci when combined with RO3306 treatment.

Based on these findings, we then considered that RO3306 treatment may activate RAD52-dependent repair. Namely, we posited that RO3306 treated cells may be more reliant on RAD52, which we assessed with clonogenic survival after 48-hour treatment with 1µM or 3µM RO3306. In cells treated with either concentration of RO3306, we observed that clonogenic survival was indeed significantly decreased in RAD52^KO^ cells as compared to RAD52^WT^ cells (siCTRL-treated), with the 3 µM RO3306 treatment having the greatest impact (Fig 4g). Next, we assessed whether siERCC6L treatment would have a similar impact on clonogenic survival in RO3306-treated cells. Beginning with RAD52^WT^ cells, 3µM RO3306 treatment (but not 1µM RO3306 treatment), induced a significant decrease in clonogenic survival when combined with siERCC6L treatment (Fig 4g). With RAD52^KO^ cells, treatment with siERCC6L caused a further decrease in viability when combined with both 1µM and 3µM RO3306 exposure (Fig 4g). Altogether, these findings indicate that ERCC6L depletion induces RAD52-GFP foci in prometaphase irrespective of CDK1 inhibition (i.e., RO3306 treatment), and that cells also rely more heavily on RAD52 when CDK1 is inhibited, which causes G2/M arrest.

### RAD52-GFP foci induced by ERCC6L depletion are distinct from sites of mitotic DNA synthesis (MiDAS)

RAD52 has been proposed to promote Mitotic DNA Synthesis (MiDAS), which is typically induced by extended treatment of cells with the DNA polymerase α inhibitor Aphidicolin (APH) (Özer and Hickson 2018; Wu 2019). Thus, we sought to define the relationship between RAD52-GFP foci and sites of MiDAS under various conditions. In particular, we compared effects of siERCC6L vs. APH treatment on both RAD52-GFP foci and induction of MiDAS. The assay for MiDAS involves detecting incorporation of the nucleoside analogue 5-ethynyl-2′-deoxyuridine (EdU) during mitosis (Garribba et al. 2018). Furthermore, MiDAS is commonly assessed using release from RO3306 treatment (Garribba et al. 2018; Bhowmick et al. 2016; Özer et al. 2018), which we also included in this analysis. Specifically, we treated cells with siERCC6L or siCTRL (all cells without siERCC6L were treated with siCTRL), then exposed cells to 0.4 µM APH for 46 hours, synchronized cells with RO3306 in the last 6 hours of APH exposure, released cells into media containing EdU for 30 minutes prior to fixation, and analyzed prometaphase cells by IF to detect EdU and RAD52-GFP (Fig 5a-b). From this analysis, we found that APH treatment induced significant increases in RAD52-GFP foci (Fig 5c). In addition, siERCC6L treatment also caused an increase in RAD52-GFP foci (Fig 5c), similar to results described above for prometaphase cells without APH treatment (Fig 4c, f). However, combining APH and siERCC6L treatment did not induce a significant increase vs. APH-treatment alone (Fig 5b-c). We next quantified EdU foci and found that APH treatment produced significantly higher levels of EdU foci compared to untreated cells, but that siERCC6L treatment did not cause an obvious increase in EdU foci, both with and without APH treatment (Fig 5b,d). We also analyzed these data to specifically examine cells with high levels of MiDAS (i.e., 5 or more EdU foci in prometaphase cells after APH treatment) and found no obvious effect of siERCC6L treatment (Fig 5e). For comparison, we also examined HU treatment conditions that caused RAD52-GFP foci (Supplementary Fig 3a) and found that such treatment did not produce a significant increase in EdU foci as compared to untreated cells, regardless of siERCC6L treatment (Supplementary Fig 3b). In summary, while both APH treatment and siERCC6L treatment cause RAD52-GFP foci in prometaphase cells, the combined treatment of APH and siERCC6L did not produce higher levels of RAD52-GFP foci vs. APH alone, and only APH treatment caused an increase in MiDAS.

**Figure 5:**
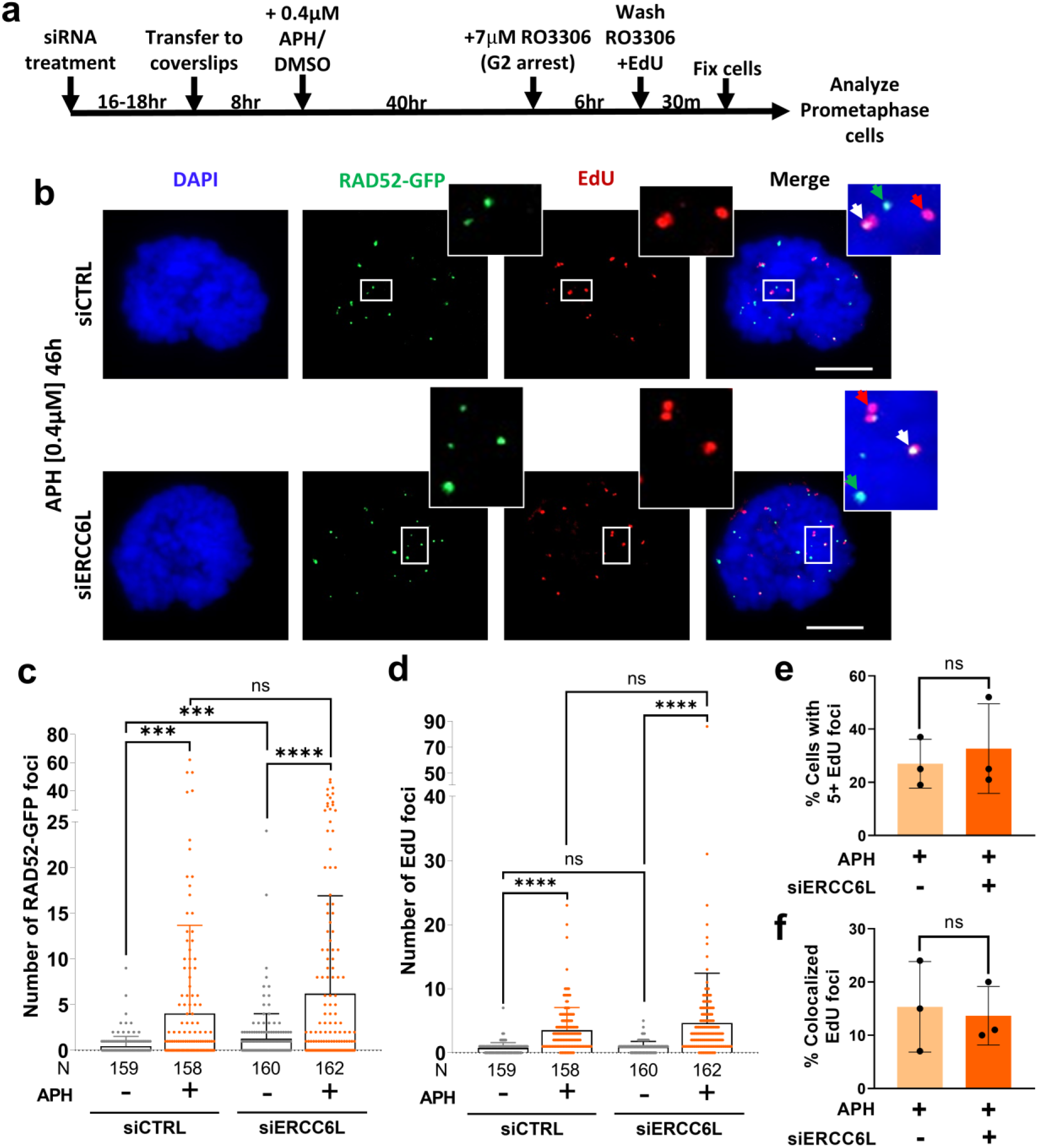
RAD52 foci induced by ERCC6L depletion are distinct from sites of mitotic DNA synthesis (MiDAS). **a**a) Schematic of treatments used for MiDAS detection in prometaphase cells expressing RAD52-GFP. **b)** Shown are representative images of cells with RAD52-GFP and EdU foci. Merge and zoom images show individual RAD52-GFP (green arrow), EdU (red arrow), and colocalized foci (white arrow). Scale bars are 10µm, and images were taken at 40x magnification. **c)** RAD52-GFP foci are induced by treatments with aphidicolin (APH, 46 hours) and siERCC6L, but are not further increased with combined treatment of siERCC6L and APH. Treatments were performed as in (a). Bars show mean foci value. N (number of cells) is shown below each bar. ns=not significant, ***=p<0.001, and ****=p<0.0001, K-S test. **d)** EdU foci increase with APH treatment, but not with siERCC6L treatment. Treatments were as in (a). Bars show mean foci value. N is shown below each bar. ns=not significant, ****=p<0.0001, K-S test. **e)** No significant difference in the percentage of cells with 5 or more EdU foci between siCTRL and siERCC6L treated cells after APH exposure. Analysis of the data shown in (c) and (d). N=3 independent experiments. ns=not significant, unpaired *t*-test. **f)** The percentage of EdU foci that colocalize with RAD52-GFP foci in cells that have 5 or more EdU foci is not significantly different in cells treated with siCTRL or siERCC6L after APH exposure. Analysis of the data shown in (c) and (d). N=3 independent experiments. ns=not significant, unpaired *t*-test.

In addition, we tested whether EdU foci colocalized with RAD52-GFP foci, and whether such colocalization was affected by siERCC6L treatment. Specifically, we used APH-treated cells with high levels of MiDAS and examined computed distances between foci maxima, where we considered two maxima to be colocalized if they were less than 3 pixels (∼0.34µm) apart. For cells treated with siCTRL, we found that RAD52-GFP foci co-localized with MiDAS (EdU foci) relatively infrequently (15%) (Fig 5b, f). Furthermore, we found that there was no significant difference between siERCC6L-treated cells and siCTRL for such co-localization (Fig 5b, f). Altogether, these data indicate that although RAD52 localizes in prometaphase cells in response to ERCC6L depletion, HU and APH exposure, this localization is not often associated with EdU incorporation (sites of DNA synthesis). Thus, it is unlikely that MiDAS is the main compensatory mechanism in which RAD52 functions to mitigate genotoxic stress induced by ERCC6L depletion or replication stress.

### Replication Stress induces delayed RAD52-GFP foci in interphase cells with ERCC6L depletion

To understand how RAD52 might further mitigate genotoxic stress outside of mitosis, we next asked if ERCC6L depletion might also alter RAD52 localization in interphase cells. Specifically, we hypothesized that in addition to RAD52 recruitment in mitosis in response to ERCC6L depletion, unresolved replication stress in the subsequent interphase associated with ERCC6L depletion may also recruit RAD52. To test this, we treated our RAD52-GFP cell line with siERCC6L and examined RAD52-GFP foci in interphase cells (Fig 6a, DMSO treatment). We observed a significant increase in Rad52-GFP foci after siERCC6L treatment at either of two time points (i.e., fixation 48 hours and 64 hours after transfection, Fig 6b-c, and 6d, respectively, samples without HU). This result indicates that ERCC6L depletion causes RAD52-GFP foci in interphase. Given the key role that ERCC6L plays in mitosis, we suggest that interphase RAD52-GFP foci are caused by defects that are transmitted from the prior mitosis, although it is possible that ERCC6L is also playing a role in interphase to suppress these RAD52-GFP foci.

**Figure 6:**
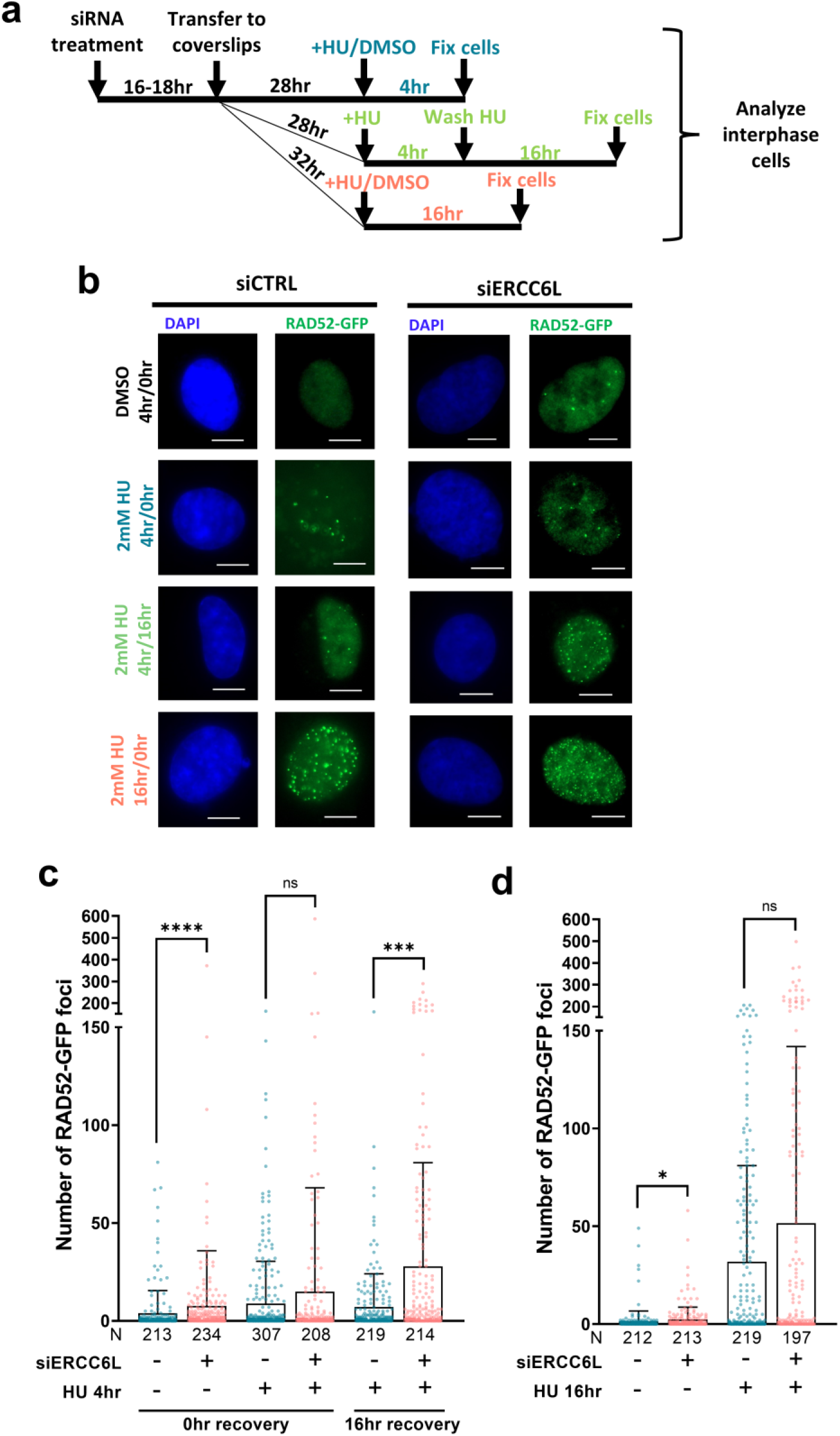
Replication Stress induces delayed RAD52-GFP foci in interphase cells with ERCC6L depletion. (**a**) Schematic of treatments to examine RAD52-GFP foci in interphase cells. **(b)** Shown are representative images of interphase cells to detect RAD52-GFP foci. Treatments were performed as shown in (a). Scale bars are 10µm and images were taken at 40x magnification. **(C)** RAD52 foci in interphase cells significantly increase with siERCC6L treatment both without HU treatment, and when combined with a 4 hr HU treatment followed by recovery for 16 hr. Bars show mean foci value. N (number of cells) is shown below each bar. ns=not significant, ***=p<0.001, and ****=p<0.0001, K-S test. **(d)** RAD52 foci in interphase cells are significantly increased by 16-hour HU exposure, which is not further increased with siERCC6L treatment. Bars show mean foci value. N is shown below each bar. ns=not significant, ***=p<0.001, and ****=p<0.0001, K-S test.

Next, we sought to examine how replication stress timing and recovery from replication stress might affect the emergence of interphase RAD52-GFP foci. Using a series of HU treatments that varied in length, and with or without recovery periods, we measured RAD52-GFP foci in response to siERCC6L or siCTRL treatment (Fig 6a-b). In cells treated with HU for 4 hours, but that were allowed to recover for 16 hours (16hr recovery), we observed significantly increased levels of RAD52-GFP foci with siERCC6L treatment (Fig 6c). By contrast, cells that were treated with HU for 4 hours without recovery (0hr recovery), or 16 hours without recovery, showed no significant difference in RAD52-GFP foci between siERCC6L and siCTRL treatments (Fig 6c, 6d, respectively). Finally, as a control, to ensure that siERCC6L treatment was not simply altering the proportion of cells in S-phase as compared to siCTRL, we stained cells with Cyclin A to assess the percentage of cells in S/G2-phase for each treatment condition (Supplementary Fig 4a). We observed that siERCC6L treatment did not substantially change the proportion of Cyclin A positive cells (S/G2-phase cells) under any HU treatment condition (Supplementary Fig 4b). In summary, the combined effect of HU treatment and siERCC6L treatment on RAD52-GFP foci required recovery from the HU treatment. We suggest that this requirement for a long recovery time to produce interphase RAD52-GFP foci reflects genotoxic stress that occurred in the prior cell cycle.

## DISCUSSION

To investigate factors and pathways important for cellular fitness in cells without RAD52, we performed genome-wide CRISPR knock-out screens, which revealed hundreds of factors with a differential effect on RAD52^KO^ vs. RAD52^WT^ cells. Using this list of genes, we performed pathway analysis to identify a series of factors for secondary screening from pathways related to the nucleus. From such secondary screening, we found that siRNA targeting of ERCC6L (also known as PICH) in RAD52^KO^ cells induces a significant increase in G1-phase 53BP1 foci formation and negatively impacts clonogenic survival. Our subsequent cell biology analysis indicates that these two factors co-compensate, in that loss of RAD52 causes elevated ERCC6L-UFBs, and depletion of ERCC6L causes elevated RAD52-GFP foci in prometaphase and interphase cells. Based on these findings, we propose the following model. RAD52 is important to mitigate replication stress, and this role is amplified in the absence of ERCC6L (Fig 7a). In the absence of RAD52, replication stress persists into mitosis where lesions are converted into DNA UFBs in mitotic anaphase. These UFBs can be resolved through a process that requires ERCC6L to avoid further accumulation of genotoxic damage and promote cell survival (Fig 7b). Likewise, loss of ERCC6L disrupts the resolution of UFBs, thereby promoting genotoxic stress that leads to RAD52-GFP foci, but combined with loss RAD52, such stress persists and causes loss of viability (Fig 7c).

**Figure 7:**
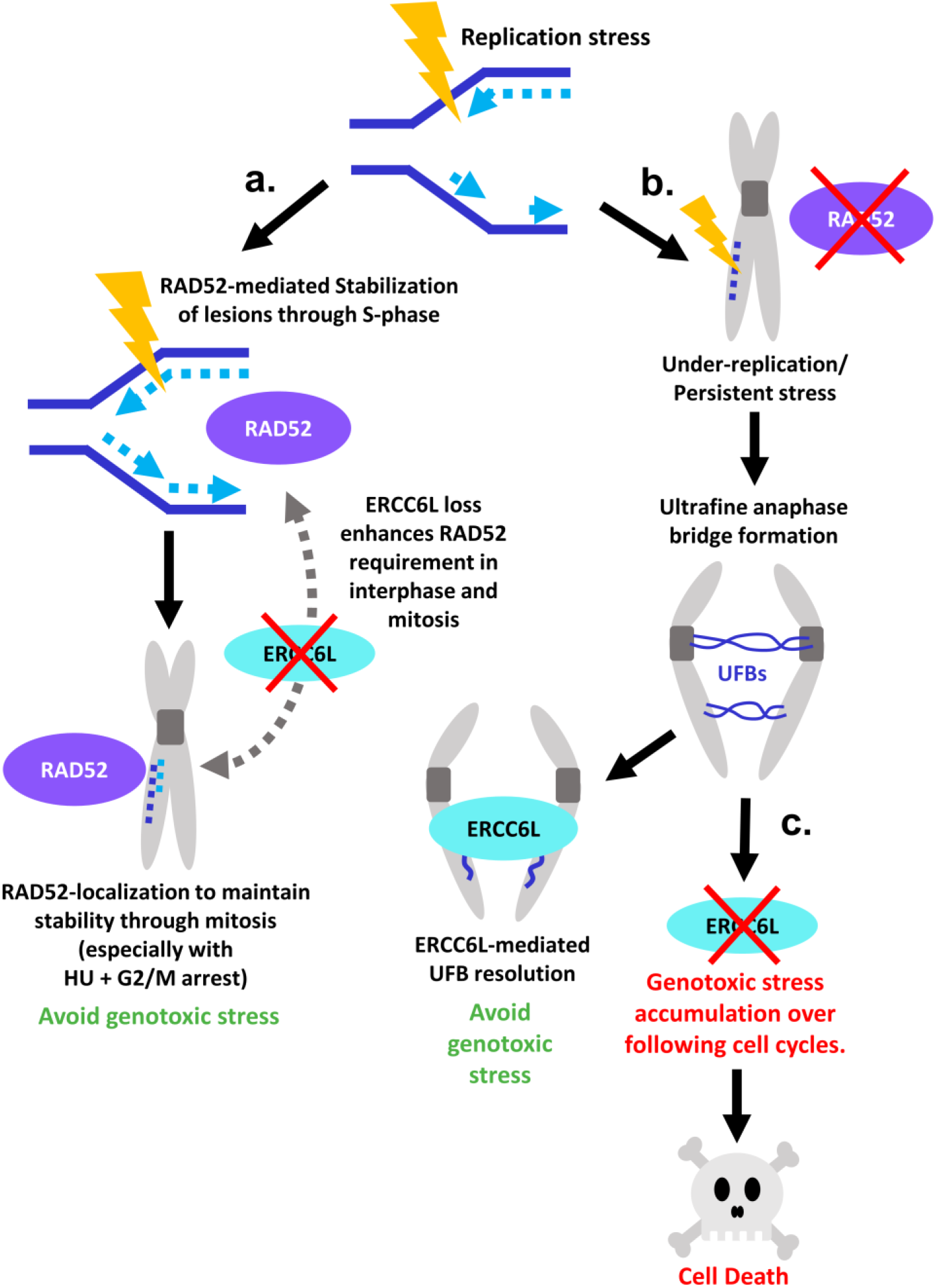
Proposed mechanism of the RAD52 ERCC6L synthetic lethal relationship. (**a**) RAD52 maintains DNA stability and prevents later UFB formation. RAD52 becomes more essential in cells lacking ERCC6L and under increased replication stress conditions combined with delayed mitotic entry. **(b)** Without RAD52, replication stress is converted to UFBs and ERCC6L becomes essential to their resolution. **(c)** In cells with neither RAD52 nor ERCC6L, genotoxic stress is allowed to accumulate, potentially over multiple cell cycles. This cycle ultimately leads to cell death.

We observed these co-compensation effects of RAD52-GFP foci and ERCC6L-UFBs without exogenous stress, but we also examined these effects following various stress treatments. Such treatments provide insight into the types of stress that amplify the co-compensation of ERCC6L and RAD52, as well as the relative timing of these effects following the treatments. First, we examined effects of the Topo IIα inhibitor ICRF-193, which appears to cause replication stress that leads to UFBs (Gemble et al. 2020). Indeed, we found that ICRF-193 treatment induces ERCC6L-UFBs, but notably, the frequency of such UFBs was higher in the RAD52^KO^ cell line, as was also the case under unstressed conditions. We suggest that RAD52 and Topo IIα are each important to suppress replication stress that results in ERCC6L-UFBs. Alternatively, RAD52 and/or Topo IIα may promote alternative UFB resolution pathways independent of ERCC6L. In any case, we then posited that combined loss of RAD52 and ICRF-193 treatment may cause an increased reliance on ERCC6L. Consistent with this notion, we found that combined loss of RAD52, depletion of ERCC6L, and ICRF-193 treatment caused a marked loss of clonogenic survival. Altogether, these findings indicate that loss of RAD52 causes an increased reliance on ERCC6L both under unstressed conditions, and with inhibition of Topo IIα.

To examine RAD52-GFP foci formation with combinations of ERCC6L depletion and replication stress, we performed a series of HU treatments. These experiments provided an indication of when RAD52 recruitment occurs with respect to the timing of the source of genotoxic stress. In particular, we found that cells depleted of ERCC6L and exposed to HU (4 hours) with a long recovery time (16 hours) had dramatically increased RAD52-GFP foci in interphase. In contrast, with HU treatment but without a recovery time, ERCC6L depletion did not cause an increase in RAD52 interphase cells. We suggest that the specific increase in RAD52-GFP foci in interphase following ERCC6L depletion and HU treatment with a long recovery time reflects RAD52 recruitment to genotoxic stress that originated in the prior cell cycle. As for what role RAD52 is playing in interphase in response to ERCC6L depletion, one possibility is that RAD52 is needed to protect lesions resulting from unresolved UFBs from triggering further instability in the following S-phase. Recent studies have identified protective roles for RAD52 during S-phase replication, where loss of RAD52 was associated with unscheduled fork reversal and degradation, accumulation of single stranded DNA (ssDNA), and reliance on enzymes associated with error-prone processes like HR (RAD51) and origin re-priming (Polα) to recover damaged forks and mitigate further instability (Malacaria et al. 2019; Biagi et al. 2023). Thus, we suggest that RAD52 may help stabilize lesions in S-phase caused by loss of ERCC6L to prevent further genotoxic stress accumulation and genome instability.

We also examined the role of RAD52 and ERCC6L in response to another cellular stress: G2/M arrest via RO3006 treatment. Such treatment is often used to enrich for prometaphase cells, such as for MiDAS analysis (see below), but recent studies have also indicated that RO3306 alters the cellular response to replication stress, which may reflect defects in modulating the timing of G2/M progression with DNA replication (Brison et al. 2023). While we found that ERCC6L depletion caused elevated RAD52-GFP foci in prometaphase cells both with and without RO3306 treatment, HU treatment caused an increase in RAD52-GFP foci only if the HU treatment was followed by RO3306 treatment. These findings indicate that RO3306 treatment may cause an increased dependency on RAD52 in prometaphase cells, which we then assessed using clonogenic survival experiments. Indeed, we found that a long (48-hour) RO3306 exposure has a negative impact on clonogenic survival in RAD52^KO^ cells. Consistent with ERCC6L and RO3306 having distinct effects on the requirement for RAD52, we further found that the combination depletion of ERCC6L and RO3306 treatment caused a marked loss in viability in RAD52^KO^ cells. Altogether, given the role of ERCC6L in mitosis, and that RO3306 disrupts G2/M progression, these findings support a key role of RAD52 to respond to stress in mitosis.

As discussed above, our data show that RAD52-GFP forms foci in prometaphase in response to ERCC6L depletion. Initially we hypothesized that such Rad52-GFP foci were sites of mitotic DNA synthesis, since RAD52 has been implicated in such mitotic DNA synthesis (i.e., MiDAS) (Bhowmick et al. 2016; Min et al. 2017, 2019; Özer et al. 2018; Epum and Haber 2021). However, a recent study found that RAD52 is dispensable for MiDAS induced by APH in RPE-1 cells (Graber-Feesl et al. 2019). Our findings also indicate that the role of RAD52 in mitosis is not necessarily linked to MiDAS, Namely, we found that RAD52-GFP foci do not often localize with EdU incorporation, and siERCC6L treatment causes an increase in RAD52-GFP foci without causing an increase EdU foci. Thus, we propose that RAD52 may be playing a structural role in prometaphase cells to stabilize lesions related to replication stress rather than mediating DNA synthesis per se.

Notably, our initial genome-wide screens did not identify several genes that were previously confirmed to be synthetic lethal with RAD52, including BRCA1, BRCA2, and PALB2 (Lok et al. 2013; Feng et al. 2011). One possible explanation is that many of the sgRNAs for these genes in the genome-wide library used in our study did not produce a knock-out phenotype. Despite this possible pitfall of the screening technique, our screens did identify two other genes found to be synthetic lethal with RAD52 independently in other studies, RAD51D (Chun et al. 2013) and XAB2 (Sharma et al. 2021), though they were not selected as hits in our sub-screen based on the G1 53BP1 foci assay (Supplementary Fig 1, Supplementary Table 4).

Regarding potential clinical significance of these findings, recent work has shown that ERCC6L is often over-expressed in breast cancers and that knockdown of ERCC6L inhibits proliferation in breast cancer cells (Liu et al. 2018). Similarly, loss of ERCC6L in triple negative breast cancer causes chromosome instability and cell death (Huang et al. 2019). More broadly, pan-cancer studies of altered ERCC6L expression have identified increased ERCC6L expression as a marker of poor prognosis in multiple types of cancer (Lu et al. 2022). Finally, RAD52 inhibitors are in pre-clinical development (Hengel et al. 2017). Thus, we suggest that evaluating ERCC6L levels and mutations in cancer should be investigated as a potential biomarker for response to targeting RAD52, and vice versa.

## MATERIALS & METHODS

### Cell lines

The parental cell line used in these studies (referred to as RAD52^WT^) is a line of hTERT RPE-1 cells that are p53^KO^ and stably express Cas9-FLAG that was generously provided by Dr. Daniel Durocher (Lunenfeld-Tanenbaum Research Institute, University of Toronto), and cultured as described (Olivieri and Durocher 2021; Olivieri et al. 2020). The RAD52^KO^ cell line was generated from the RAD52^WT^ line by CRISPR-Cas9-mediated deletion of the 1.1kb region between exons 3 and 4 using guide RNAs sg1 and sg2 cloned into px330 (Addgene #42230) as previously described (Kelso et al. 2019). These sgRNA vectors were co-transfected with a dsRED expression vector using ViaFect transfection reagent (Promega #E4981). After 3 days, cells were sorted to enrich dsRED-positive cells (BD Aria) and plated at low density to achieve single colonies. The RAD52^KO^ cell line was then isolated through screening individual clones by PCR (oligos ol1 and ol2) and Sanger sequencing. Knock-out of the RAD52 protein was confirmed by immunoblot analysis. All sgRNAs and oligos are listed in Supplementary Table 1.

The RAD52-GFP cell line was generated by transfecting the RAD52^WT^ cell line with 800 ng of a RAD52 expression vector that is tagged with GFP at the N-terminus that was generously provided by Dr. Markus Lobrich (Technical University of Darmstadt) (Llorens-Agost et al. 2021). The transfection was performed in a 6 well with 2 mL of antibiotic free media, and using Viafect transfection reagent. Subsequently, cells were sorted to enrich for GFP-positive cells, which were cultured and sorted a second time (BD Aria). Cells were routinely checked for GFP-positive levels and periodically resorted. All cell lines used in experiments tested negative for mycoplasma contamination (Lonza MycoAlert PLUS Mycoplasma Detection Kit).

### CRISPR-KO screens

CRISPR-KO screens were carried out as described (Olivieri and Durocher 2021), but with some modifications. Specifically, RAD52^KO^ and RAD52^WT^ cells were transduced at a low multiplicity of infection (MOI<0.3) using the BFP-tagged Human Improved Genome-wide Knockout CRIPSR Library (Addgene #67989) (Tzelepis et al. 2016) packaged into lentivirus (via 293T cells). MOI was confirmed for both cell lines two days after transduction by flow cytometry analysis (Quanteon) where MOI was equivalent to the percentage of BFP-positive cells. Following transduction, cells were treated with media containing 20 µg/ml puromycin to select for and enrich cells transduced with the library. After 24 hours, cells were trypsinized and resuspended in fresh media with 20 µg/ml puromycin and incubated for another 24 hours. At 2 days post-transduction (considered to be the initial time point, T0), puromycin was removed. At 8 days post-transduction (T6), the total cells were split into 3 different treatment groups: Untreated, 2 Gy Ionizing Radiation exposure (IR), and 1 µM cisplatin treatment (Cis-Pt). Cells in the IR screen were irradiated once (Gammacell 3000) on T6, while cells in the Cis-Pt screen were grown in media containing 1 µM cisplatin for 6 days (T6-T12). Cells were passaged to maintain 400x coverage of the CRISPR library every 3 days through the duration of the screening period (T3-T18).

Cell pellets were collected from T0 and T18 time points for analysis, where the number of cells in each pellet (∼36 million cells) was set to represent the total number of guides in the CRISPR library (90,709 guides) multiplied by the desired library coverage for sequencing (400x). Cell pellets were processed into genomic DNA using the QIAmp Blood Maxi Kit (Qiagen #51192), and the integrated sgRNA library was amplified from the genomic DNA by PCR using Q5 Hot Start High-Fidelity 2x Master Mix (NEB #M0494L) and oligos ol3 and ol4 (Supplementary Table 1). Amplified and gel purified PCR products were sequenced by Illumina HiSeq (Azenta), and the resulting reads were analyzed using the maximum-likelihood estimation (MLE) algorithm from the MAGeCK pipeline(Li et al. 2014) to generate sgRNA counts and selectivity (Beta) scores comparing the RAD52^WT^ and RAD52^KO^ cell lines in each of the three screens.

### Gene hit selection for secondary screening

The MAGeCKFlute pipeline (Wang et al. 2019) was used to perform downstream analyses for results from MAGeCK MLE results, described above. This pipeline includes normalization of beta scores to account for cell division rate differences, and determination of gene hits by applying standard deviation cutoffs to plotted normalized Beta scores of 1.5x the standard deviation to the x, y, and diagonal axes. Gene “hits” were selected as those in the section of the plot that were within the x-axis cutoffs (RAD52^WT^ beta score 1.5x standard deviation from x=0), but beyond the negative y-axis and diagonal cutoffs (RAD52^KO^ beta score <1.5x standard deviation from y=0 and diagonal) (see Fig 1c and Supplementary Table 2). These gene hits from all three screens were combined for gene set enrichment analysis (GSEA) with MAGeCKFlute using default parameters. GSEA enriched pathways with FDR <0.01 and NES scores of >0.5 from all three screen conditions were categorized into 6 groups (Fig 1e) based on the enriched pathway names. Using the three groups with pathways functioning inside the nucleus, at least one gene from each enriched pathway was selected, and genes that were reported as hits in multiple pathways, multiple screen conditions, or multiple components of the same complex were favored.

### Immunoblot analysis

Cells were lysed with either NETN buffer (20 mM TRIS (pH 8.0), 100 mM NaCl, 1 mM ethylenediaminetetraacetic acid (EDTA), 0.5% IGEPAL, 1.25 mM dithiothreitol and Roche Protease Inhibitor) and a series of freeze-thaw cycles (RAD52 blot), or ELB buffer (250 mM NaCI, 5 mM EDTA, 50 mM HEPES, 0.1% Ipegal, Roche protease inhibitor) with sonication (Qsonica, #Q800R) (Cas9-FLAG and ERCC6L blots). Blots were probed using antibodies against RAD52 1:500 (Santa Cruz Biotechnology #sc365341), FLAG 1:1000 (Sigma #A8592), ERCC6L 1:500 (Abnova #H00054821-D01P), or ACTIN 1:1000 (Sigma #A2066), and with HRP-conjugated secondary antibodies: rabbit anti-mouse 1:3000 (Abcam #ab205719) or goat anti-rabbit 1:3000 (Abcam #ab205718). ECL Western Blotting Substrate (Thermo Fisher Scientific #32106) was used to detect HRP signal on film.

### Cell biology assays

For the G1 53BP1 foci assay, RAD52^KO^ and RAD52^WT^ cells were seeded at 5 x 10^4^ cells per well in 12-well plates with 1 mL of antibiotic-free media and treated with Lipofectamine RNAiMAX (ThermoFisher #13778075) and 10 pmol of siRNA pools (pool of 4 siRNAs; Dharmacon siGENOME, sequences from manufacturer in Supplementary Table 1) or non-targeting siRNA (siCTRL; Dharmacon #D-001810-01-20). After 16-18 hours, cells were trypsinized, resuspended, and transferred to coverslips in 6-well plates. 3 Days after siRNA treatment, cells were fixed in 4% paraformaldehyde (PFA), permeabilized with 0.5% Triton X-100 in PBS, and blocked with 8% goat serum in PBS. Fixed cells were incubated with rabbit 53BP1 antibody 1:500 (Abcam #ab36823) and mouse Cyclin A antibody 1:100 (Santa Cruz #sc-271682). Secondary antibodies (goat-anti-rabbit 488 (Invitrogen #A-32731) and goat-anti-mouse 594 (Invitrogen #A-11032) were applied at 1:500.

For the Anaphase ultra-fine bridge assay, RAD52^KO^ and RAD52^WT^ cells were seeded at 1 x 10^5^ cells per well in 6-well plates with 2 mL of antibiotic-free media along with Lipofectamine RNAiMAX and 20 pmol of siCTRL. After 16-18 hours, cells were trypsinized, resuspended, and transferred to coverslips in 6-well plates. For experiments where cells were treated with ICRF-193 (Enzo Life Sciences #BML-GR332-0001), 3 days after transfection, 200 nM ICRF-193 or equivalent volume of vehicle (DMSO) was added to each well for 6 hours prior to fixation. Afterwards, cells were washed and allowed to recover for 16 hours prior to fixation. For all experiments, cells were pre-extracted with buffer A (0.2% Triton X-100 in 1x PEM buffer: 10x PEM buffer: 200mM HEPES pH 7.2-7.4, 10mM MgCl_2_, 100mM EGTA, diluted to 1x in PBS), and fixed in buffer B (0.1% Triton X-100, 8% PFA in 1x PEM buffer), which is similar to previously described(Bizard et al. 2018). Fixed cells were blocked in PGST (8% goat serum, 0.5% Triton-X-100 in PBS) for at least 24 hours at 4°C. Cells were then incubated at 4°C overnight with rabbit ERCC6L antibody (Abnova #H00054821-D01P, diluted 1:100) prepared on ice. Cells were then incubated with goat-anti-rabbit 488 (Invitrogen) secondary antibody.

For RAD52-GFP foci experiments, RAD52-GFP-expressing cells were seeded at 1 x 10^5^ cells per well in 6-well plates with 2 mL of antibiotic-free media along with Lipofectamine RNAiMAX and 20 pmol of either a pool of 4 siRNAs targeting ERCC6L (siERCC6L; Dharmacon #M-031581-01-0005, sequences from manufacturer in Supplemental Table 1) or siCTRL. After 16-18 hours, cells were trypsinized, resuspended, and transferred to coverslips in 6-well plates. For experiments assaying prometaphase cells, 2 days after transfection, cells were exposed to 2 mM HU or equivalent volume of DMSO for 4 hours, washed, and allowed to recover in fresh media for 12 hours. In experiments where cells were synchronized in G2/M and released into mitosis prior to fixation, 7 µM RO3306 (CDK1 inhibitor; Selleck chemicals #S7747) was added and cells were incubated for 6 hours. Cells were then washed, incubated for 30 minutes in fresh media, and fixed. In experiments where cells were not synchronized, cells were allowed to recover from HU exposure for 18 hours in fresh media and then fixed. For experiments assaying interphase cells with a 4-hour HU exposure, 2 mM HU or equivalent volume of DMSO was added to cells 2 days after transfection and incubated for 4 hours. Experiments without recovery time were fixed immediately after the 4-hour exposure, while experiments with recovery were washed and allowed to recover for 16 hours in fresh media prior to fixation. For experiments with 16-hour HU exposure (interphase cell experiments only), 2mM HU or equivalent volume of DMSO was added to cells 2 days after transfection and cells were incubated for 16 hours followed by fixation. For all RAD52-GFP experiments, cells were fixed by the same method used for anaphase ultrafine bridge assay, which is similar to previously described (Bizard et al. 2018).

In experiments where interphase cells were analyzed, fixed cells were blocked with PGST for at least 24 hours at 4°C. To evaluate cell cycle profiles, one set of experiments was also incubated with mouse Cyclin A antibody (1:100), followed by goat-anti-mouse 594 secondary antibody (1:500).

For the MiDAS assay, RAD52-GFP-expressing cells were seeded and transfected with siRNA similarly to RAD52-GFP foci experiments. For experiments with Aphidicolin (APH) treatment, cells were treated with 0.4µM APH or equivalent volume of DMSO (added 2 days after transfection) for a total of 46 hours. For experiments with HU treatment, 2 mM HU or equivalent volume of DMSO was added 3 days after transfection, and cells were incubated for 4 hours. HU-treated cells were washed and allowed to recover for 12 hours. Both HU– and APH-treated cells (and DMSO-treated controls) were then synchronized by 7µM RO3306 treatment for 6 hours. Cells were washed and released into mitosis in media containing 20µM 5-ethynyl-2’-deoxyuridine (EdU) and incubated for 30 minutes. Cells were fixed using the same method used for anaphase ultrafine bridge assay, which is similar to previously described(Bizard et al. 2018). Fixed cells were processed using the EdU Click-iT detection kit (Thermofisher #C10639) with modified reaction components, as previously described (Garribba et al. 2018).

For mounting, imaging acquisition, and image analysis, coverslips from all cell biology experiments were mounted with VECTASHIELD antifade mounting medium with DAPI (Vector labs #H-1200-10) prior to imaging. Images from the anaphase ultra-fine bridge assay experiments were acquired using an Olympus IX71 microscope with a 60x oil objective. Images from all other cell biology experiments were acquired using a ZEISS Axio Observer II microscope with a 20x air objective (G1 53BP1 assay) or 40x oil objective (RAD52-GFP foci, and MiDAS assays). Foci (53BP1, RAD52-GFP, and EdU) and Cyclin A intensity were quantified using Fiji (ImageJ). Anaphase ERCC6L ultra-fine bridges were quantified manually. Colocalization analysis was performed using an automated ImageJ macro to determine XY positions of foci maxima. Distances between foci were then calculated using the following formula where x and y are the X and Y coordinates for focus 1 vs. focus 2:

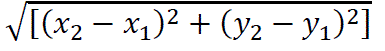

Foci were defined as colocalized if the distance between the two foci maxima was less than 3 pixels (∼0.34µm). The ImageJ macro and excel sheet used for comparison of foci maxima can be downloaded from https://github.com/baosia/FociXY.

### Clonogenic survival assays

RAD52^KO^ and RAD52^WT^ cells were seeded at 3 x 10^4^ cells per well in 12-well plates with 1 mL of antibiotic-free media and treated with Lipofectamine RNAiMAX and 10 pmol of siRNA pools (Supplemental Table 1), or siCTRL. After 16-18 hours, cells were trypsinized, resuspended, and re-seeded at low-density to allow the formation of colonies in 6-well plates. For drug-treated experiments, ICRF-193, RO3306, or equivalent volumes of DMSO were added to cells on day 3 and washed from cells 55 hours (ICRF-193) or 48 hours (RO3306) later. 11 days after cells were first seeded, cells were fixed in 10% formalin, stained with 0.5% crystal violet in 25% methanol, and counted using a 10x objective. Clonogenic survival frequencies were calculated by normalizing the number of colonies counted per well to the number of cells plated (plating efficiency) and then normalizing this value from each well to the mean of the parallel siCTRL/vehicle treated wells (siCTRL/DMSO=1).

### Author Contributions

B.O., A.M., F.W.L., and X.P. developed methods/reagents or performed the experiments. B.O. and A.M. performed data quantification and analysis. B.O. and J.M.S designed the experiments, interpreted the data, and wrote the manuscript. All authors have read and approved the submitted version of the manuscript.

---

**Supplementary Figure 1:**
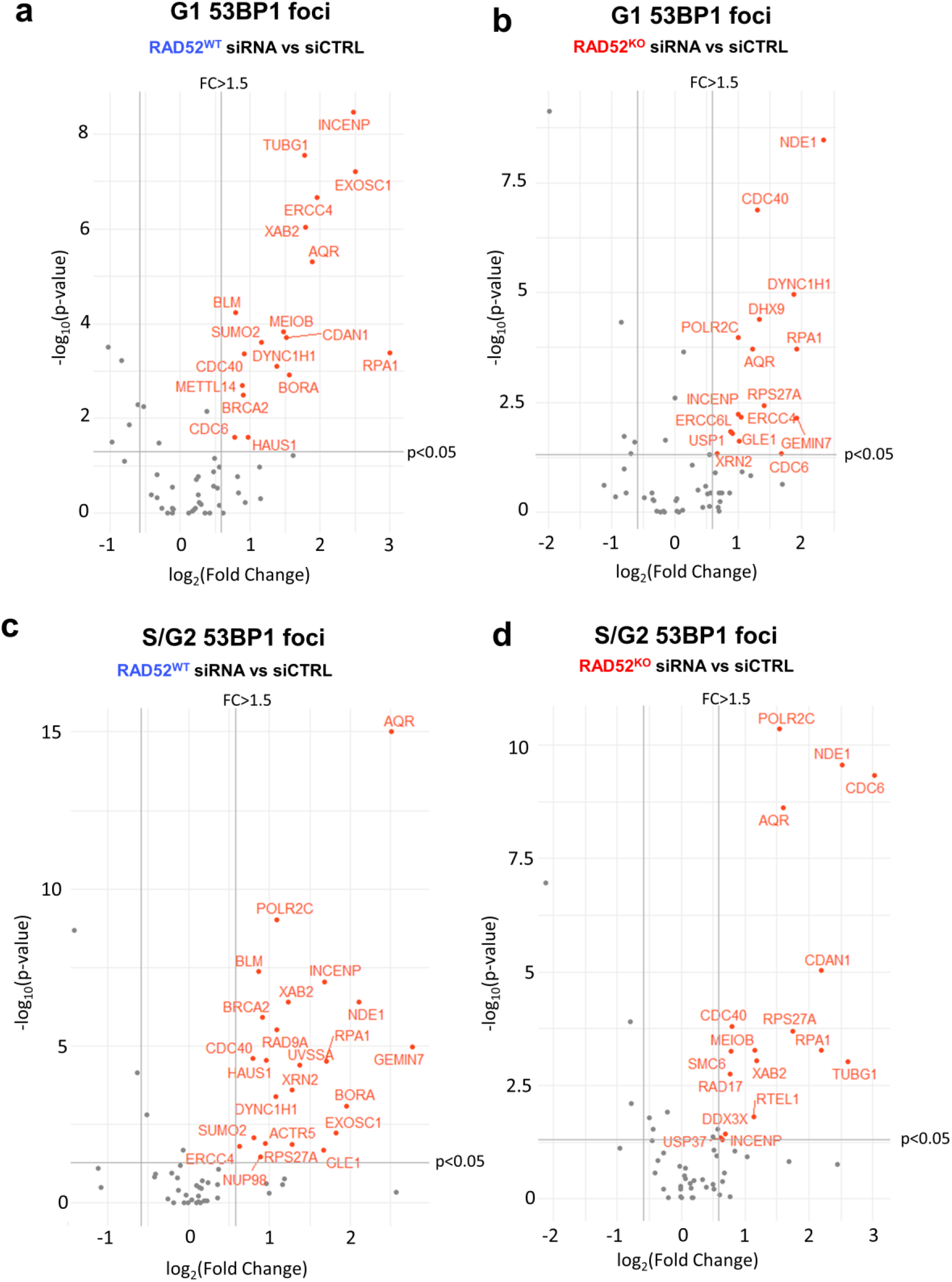
Summary of sub-screen results for 60 genes assessed for 53BP1 foci levels. siRNA sub-screen hits from IN pathway groups 1-3 (59 genes + BRCA2, see Fig 1d) that have a significant (p<0.05 by Kolmogorov-Smirnov test) and >1.5-fold mean increase in G1 53BP1 foci in the RAD52^WT^ cell line **(a)**, G1 53BP1 foci in the RAD52^KO^ cell line **(b)**, S/G2 53BP1 foci in the RAD52^WT^ cell line **(c)**, and S/G2 53BP1 foci in the RAD52^KO^ cell line **(d)** are shown in red. For all siRNAs shown, N>50 nuclei were analyzed in both RAD52^KO^ and RAD52^WT^ lines.

**Supplementary Figure 2:**
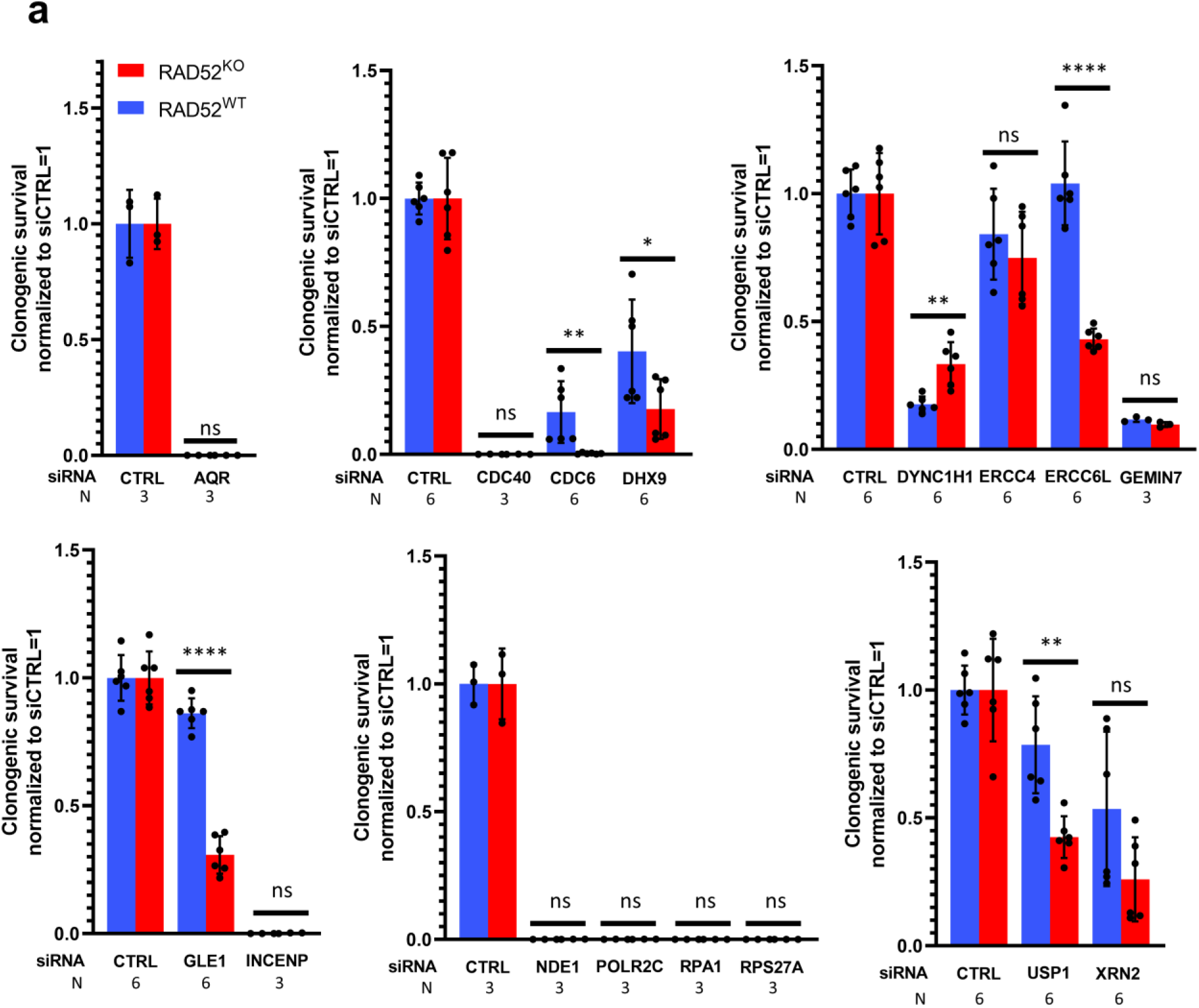
Impact of depletion of sub-screen hits on clonogenic survival. **a**) Clonogenic survival assay results for all 16 hits from the sub-screen (G1 53BP1 foci counts). Colony count data is shown for RAD52^KO^ vs RAD52^WT^ lines with depletion of the 16 hits or siCTRL treatment and are normalized to respective siCTRL treated lines (siCTRL=1). Statistical significance is determined by unpaired t-test, where ns=not significant, *=p<0.05, **=p<0.01, ***=p<0.001, ****=p<0.0001. The number (N) of replicates is listed below each set of bars and reflects the number for both the RAD52^KO^ and RAD52^WT^ lines.

**Supplementary Figure 3:**
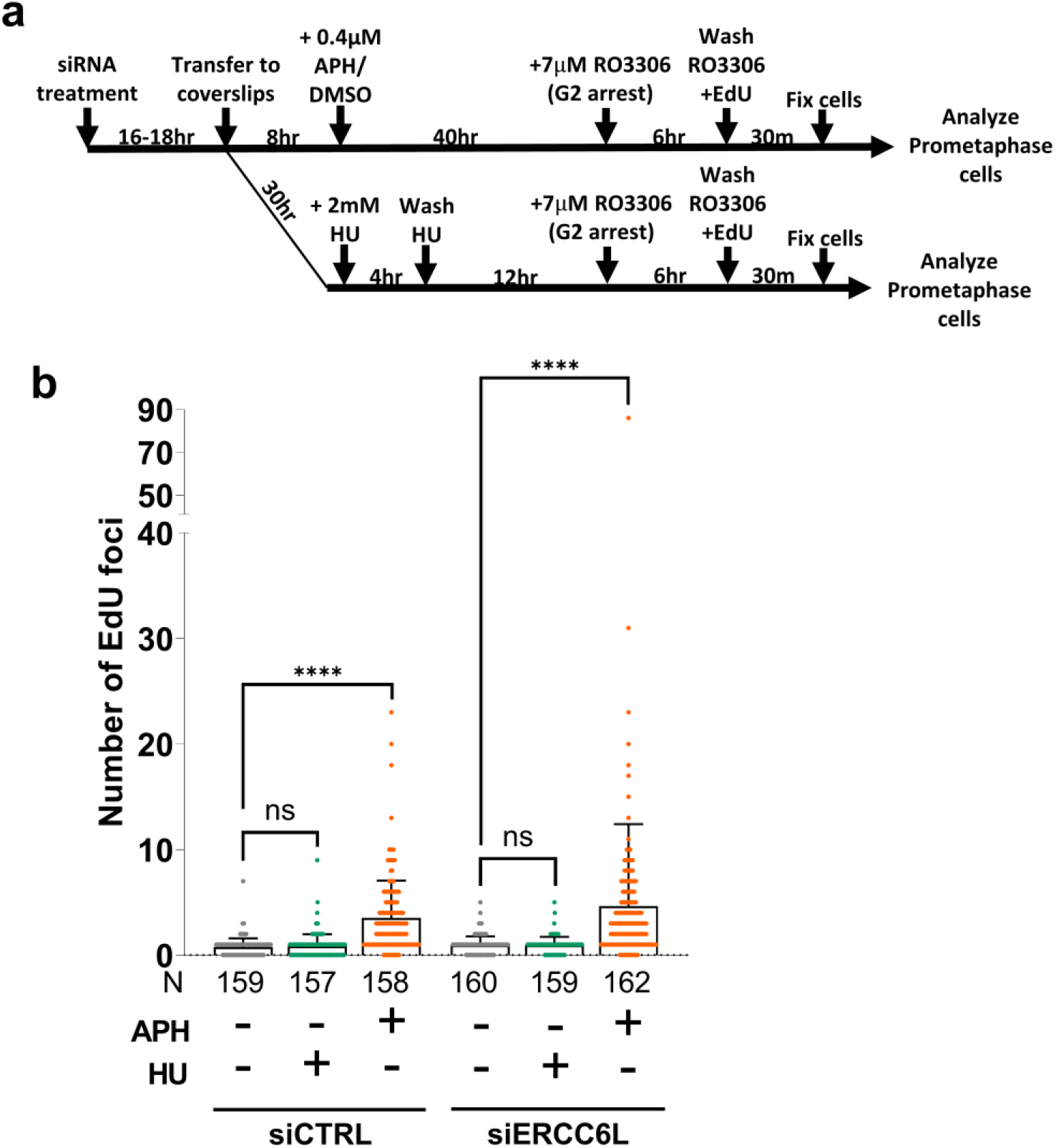
HU does not induce EdU foci in both siERCC6L– and siCTRL-treated cells. **a**) Schematic of treatments used for MiDAS detection in RAD52-GFP-expressing prometaphase cells exposed to either APH or HU. **b)** EdU foci increase upon APH exposure but not with HU exposure and recovery, irrespective of siERCC6L treatment. Bars show mean foci value. Significance determined by K-S test where ns=not significant and ****=p<0.0001.

**Supplementary Figure 4:**
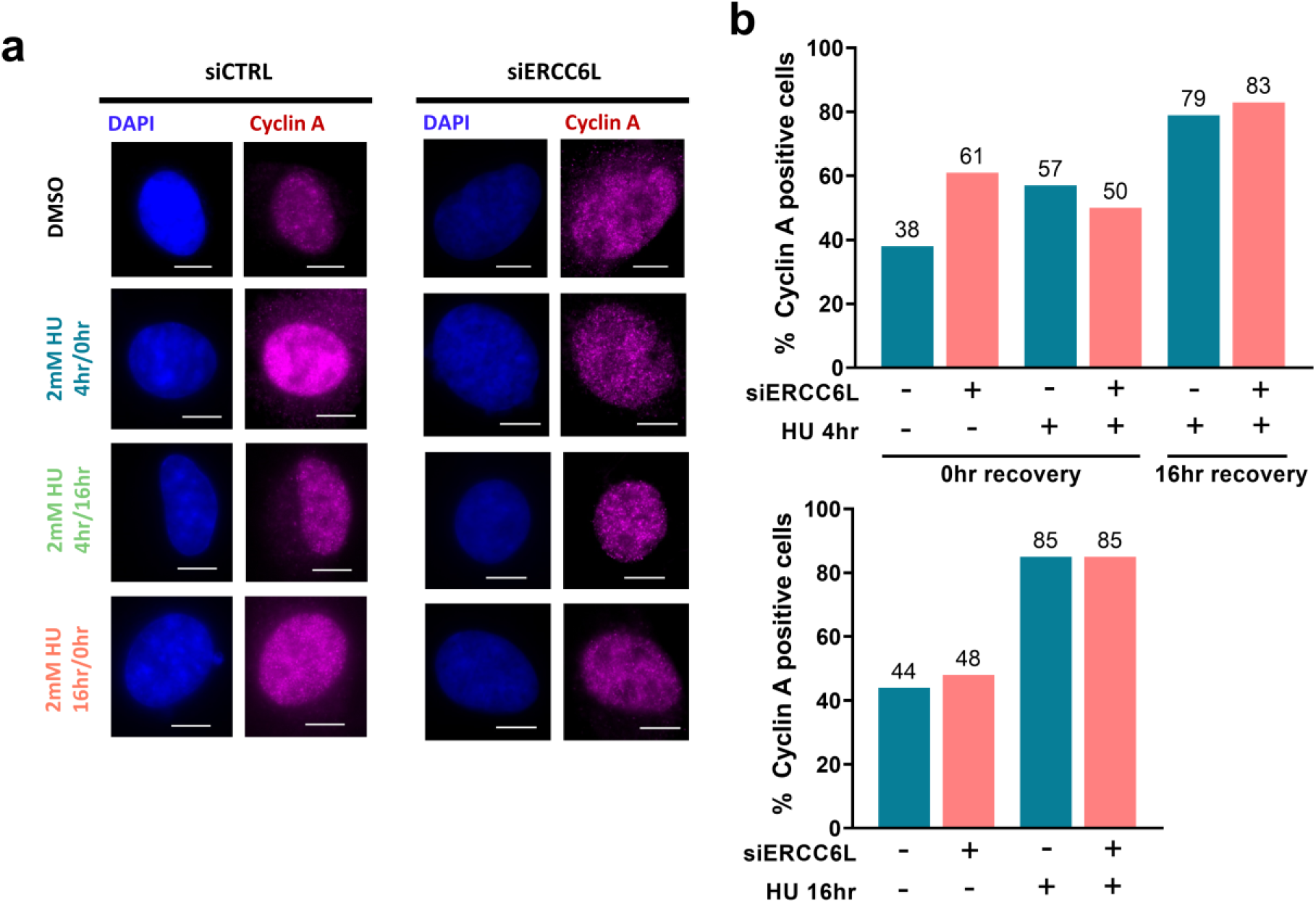
ERCC6L-depletion does not alter the percentage of Cyclin A-positive cells. **a**) Example IF images of interphase cells with Cyclin A co-staining (RAD52 foci are shown in Fig 6b). Treatments with siERCC6L/siCTRL and HU are as shown in Fig 6a. Scale bars are 10µm and images were taken at 40x magnification. **b)** siERCC6L treatment does not substantially alter Cyclin A positive levels in interphase cells. HU treatment durations and recovery times are as shown in Fig 6a,c-d.

